# The dark side of amyloid aggregation: Exploring the productive and non-productive pathways with multi-scale modeling

**DOI:** 10.1101/687020

**Authors:** Zhiguang Jia, Jeremy D. Schmit, Jianhan Chen

## Abstract

Atomistic description of protein fibril formation has remained prohibitive due to the complexity and long timescales of the conformational search problem. Here, we develop a multi-scale approach that combines a large number of atomistic molecular dynamics simulations in explicit solvent to derive Markov State Models (MSMs) for simulation of fibril growth. The search for the in-registered fully bound fibril state is modeled as a random walk on a rugged 2D energy landscape along enumerated β-sheet registry and hydrogen bonding states, whereas interconversions among nonspecific bound states and between nonspecific and hydrogen-bounded states are derived from kinetic clustering analysis. The reversible association/dissociation of an incoming peptide and overall growth kinetics are then computed from MSM trajectories. This approach is applied to derive a comprehensive description of fibril elongation of wild-type Aβ_16-22_ and how it is modulated by phenylalanine to cyclohexane (CHA) mutations. The resulting models recapitulate the experimental observation that mutants CHA19 and CHA1920 accelerate fibril elongation, but have a relatively minor effect on the critical concentration for fibril growth. Importantly, the kinetic consequences of mutations arise from a complex perturbation of the network of productive and non-productive pathways of fibril grown. This is consistent with the expectation that non-functional states will not have evolved efficient folding pathways and, therefore, will require a random search of configuration space. This study highlights the importance of describing the complete energy landscape when studying the elongation mechanism and kinetics of protein fibrils.

## Introduction

Aggregation of mis-folded proteins has been implicated in many devastating and currently incurable medical disorders such as like Alzheimer’s and prion diseases.(1-3) Originally, insoluble amyloid fibrils were considered to be the main neurotoxic agents. However, recent evidence suggests that soluble metastable amyloid β (Aβ) oligomers are more likely responsible for neuronal injury occurring in Alzheimer’s disease (AD).(4-8) This observation implies that inhibition of oligomer formation or even promotion of fibril formation could be a promising therapeutic strategy.(9, 10) However, owing to the complexity of the aggregation process and metastability of the intermediate states, identifying the most promising target species for therapeutic intervention is still challenging.(11) Therefore, it is important to deduce molecular mechanisms that control the kinetics of state transitions which, in turn, determine the populations of different Aβ aggregate species.

Elucidating the molecular mechanisms of amyloid aggregation requires detailed knowledge of the kinetics and thermodynamics of substate transitions during aggregation. Previous experiments suggest Aβ fibrils are likely grown one monomer a time, which can be described by a two-step (Dock and Lock) model.(12-15) In this model, an Aβ monomer rapidly adheres to a preformed fibril (dock step), followed by a slow lock step where the unstructured incoming peptide rearranges into an extended fibrillar conformation.(8, 12, 16) However, further mechanistic dissection of the aggregation process using experimental techniques is hindered by the presence of numerous metastable substates that interconvert rapidly.(3) Molecular dynamics (MD) simulation is a powerful tool which can provide a detailed molecular picture of the dynamics to complement experimental studies (17-19). However, the experimentally estimated elongation rate is roughly about a second per layer,(20, 21) which is beyond the currently feasible timescale with all-atom/united-atom simulations. Although this timescale is available by simplified coarse-grained (CG) approach,(19, 22-26) reduced resolution of these models makes it challenging to properly distinguish protein sidechains with similar properties and capture mutational effects on the amyloid aggregation.

The slow elongation rate of amyloid fibrils is due to a relatively flat energy landscape which is pitted with numerous local minima.(27) Since the long aggregation time arises from transitions between substates, the problem of simulating the elongation process can be solved by efficiently sampling the transitions between different substates with proper reaction coordinates. Multi-scale methods and enhanced sampling have been applied to the simulation of amyloid aggregation.(28-31) However, reproducing the kinetic network requires not only the productive pathways, but also non-productive pathways, to be adequately sampled, which is challenging for currently available enhanced-sampling techniques.

We have recently developed a theoretical and computational framework that can systematically explore productive and unproductive pathways of protein fibril growth.(27, 32) In this framework, the hydrogen bonding (H-bond) number and alignment (referred to as the ‘registry’ here) between an incoming peptide and the existing fibril core are used as the primary reaction coordinates of the aggregation process. The slow process of exploring different registries is then converted to a series of H-bond formation and breakage events. Since the H-bond formation and breakage occur on the nanosecond timescale, carefully designed simulations can be deployed to calculate the kinetic parameters of these events and measure how these parameters are affected by variations in protein sequence and/or environmental factors. These microscopic kinetic parameters can be then used to construct Markov State Models (MSMs) for simulating the overall binding and fibril growth process, This is similar to the application of MSMs for protein folding with the important difference that our state space is informed by theory (31, 33-38).

We have previously deployed this multi-scale computational algorithm to predict the binding and aggregation process of small amyloid-forming peptides in implicit solvent (27). That work established the feasibility of using the analytic theory as a framework to access long timescales and more extensive sampling of trajectories. It also successfully recapitulated the effects of single and double mutants on the aggregation kinetics of an amyloid β fragment (Aβ_16-22_)(1, 39). Specifically, the simulation reproduces the non-additive effects of introducing more hydrophobic sidechains, via Phe to cyclohexylalanine (CHA) mutation, on aggregation kinetics.(40) The order of growth rates of WT and two single mutations (WT << CHA19 < CHA20) is correctly predicted by our MSM models derived from implicit solvent simulations. The simulation also correctly predicts that the double mutation (CHA19CHA20) will reduce the fibril growth rate relative to CHA20. However, an inspection of the microscopic pathways led to an unsatisfying conclusion that the non-additive effects arose from the accumulation of small perturbations across the conformational landscape. Furthermore, the MSM had quantitative discrepancies with experiment, highlighted by an over-estimation of the deleterious effect of the double mutant.

In the present work we improve upon our earlier effort by adapting our method for use in explicit solvent. This required an expansion of the state space in the MSM to include the backbone conformational states. We also include a more careful treatment of the weakly bound states that were not considered in the analytic theory. These states (henceforth referred to as non-registered states) lack backbone H-bonds with the fibril and serve as a hub connecting the H-bonding states, which motivates our efforts to sample and include them in an expanded MSM. The improvements result in a satisfying improvement in the quantitative agreement with experiment. Perhaps more importantly, the complex effects of the mutation can be understood based on the funnel model of protein folding (41-43), which implies that non-functional states will not evolve efficient folding pathways that are dominated by a small number of trajectories. This situation requires the development of efficient algorithms, such as the one presented here, to properly sample the many small perturbations throughout conformational space that contribute to mutation outcomes.

## Results and Discussions

### MSM modeling of protein fibril growth

Our model (Fig. 1) is based on the discretization of the elongation process into a series of states based on the microscopic theory of fibril elongation.(27, 32) The elongation process starts from an unstructured monomer in the bulk solvent, which “docks” to a pre-formed fibril core and “locks” as a new structured layer.(12, 13, 16) Thus, the process can be divided into three stages. The first stage is a diffusion-controlled process, in which the incoming peptide migrates from solvent to a separation distance *b*.(44, 45) Our treatment of this stage is similar to the Brownian dynamics treatment of diffusion controlled reactions(44, 45), in which simulations starting from a monomer at the *b* surface were performed, and the rate of monomers reaching the fibril surface or the monomers reaching an escape distance *q* were analyzed. If the monomer reaches the *q* surface, the monomer is considered to have returned to the bulk solvent. The second stage describes the nonspecific interaction between the monomer and fibril without either in-register or off-register backbone H-bonds (non-registered states, blue box in Fig.1). This stage ends when the incoming peptide either forms backbone H-bonds with the core peptide, thereby entering the registered stage (third stage), or dissociates from the fibril surface, returning to the unbound state (the *q* surface). The third stage represents the peptide’s exploration of β-strand alignments by forming or breaking H-bonds (green box in Fig. 1). The attempt ends when the molecule either forms a full set of H-bonds in the correct registry (red arrow in Fig. 1, right panel) or falls off the end of the fibril (returns to the non-registered states). By combining the three stages, our model can describe the complete association and dissociation process of fibril growth. The overall growth rate can be expressed as

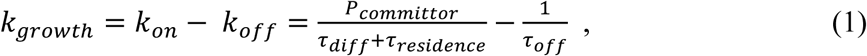

in which *τ*_*diff*_ is the average time for a monomer to diffuse from bulk solvent to the *b* surface, *τ*_*residence*_ is the average time required for a molecule entering the *b* surface to evolve to either the fully bound state or return to a separation distance *q. τ*_*off*_ represents the average time required for a fully bound peptide to reach the dissociated state (*q* surface). *P*_*committor*_ is the probability that an incoming peptide starting from the *b* surface becomes incorporated into the fibril in a fully-bound in-register state.

**Figure 1.**
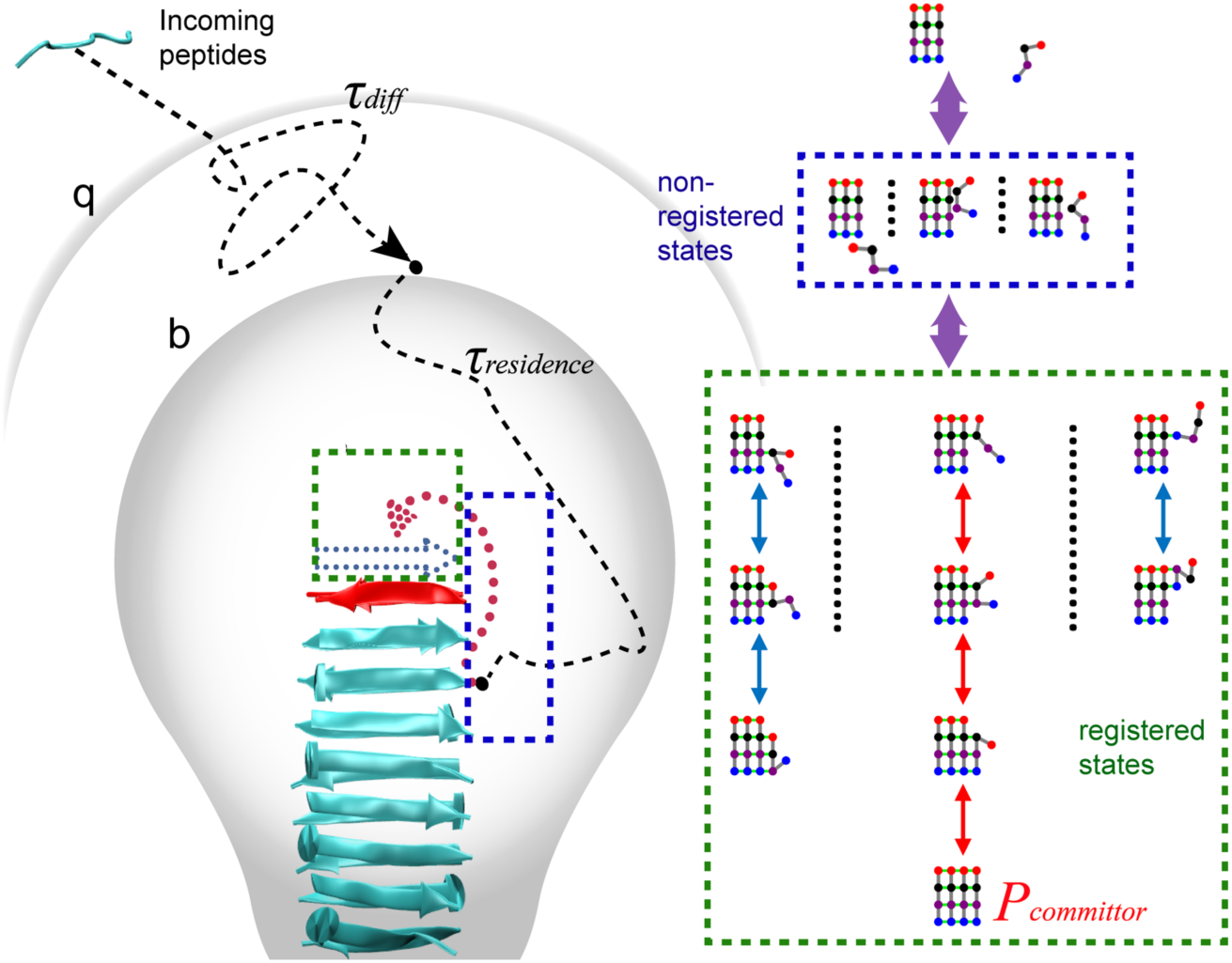
Schematic illustration of the random walk model of fibril growth with key kinetic parameters required for calculating the growth rate. The incoming peptide migrates from solvent to a separation distance *b* with a time *τ*_*diff*_ (first stage). Then within *τ*_*residence*_, it explores multiple non-registered (blue box) and registered states (green box) (second and third stage). The peptide is considered return to the bulk solvent if it reaches the *q* surface. *P*_*committor*_ is the probability of the peptide arriving at the fully-bound in-register state (the red path).

In our previous work (27), we focused on the fibril growth of the central hydrophobic core of Aβ (Aβ_16-22_, K_16_LVFFAE_22_),(46) which is among the shortest sequences that form amyloid fibrils similar to those by full-length Aβ.(46-48) All possible H-bond registry states were enumerated and MD simulations in a simple solvent accessible surface area (SASA)-based implicit solvent model (49) were performed to derive the average transition times between neighboring states. The fibril growth of the wild-type and mutant peptides (single mutation CHA19/CHA20 and double mutation CHA19CHA20) were predicted by the MSMs. However, due to the smoother energy surface of the SASA implicit solvent model and lack of solvent friction, conformational fluctuations among non-registered states of the peptide were found to be much faster than sampling of registered states, allowing a single non-registered state to be used in the MSM. Furthermore, simple implicit solvent models like SASA are not capable of accurately describing the peptide conformational equilibria(50, 51). These factors likely contribute to over-estimation of the reduction in the growth rate due to the double mutation (CHA19CHA20) compared to CHA19 and CHA20 single mutants. To overcome this, explicit solvent simulations were deployed in the current work, which enables more realistic transition rates to be computed and allows better sampling of peptide conformational fluctuations in non-registered as well as registered states. The non-registered conformations sampled will be clustered based on the peptide structure and peptide-fibril contacts. These clustered states will then be added to the MSM. In addition, we will introduce a new type of transition between registered states, which involves peptide conformational changes. These transitions will also be necessary when extending the current framework to longer peptides.

### H-bond transition rates of wild type and mutated Aβ_16-22_ peptides

We define registered states by orientation of the incoming peptide (antiparallel or parallel), surfaces of the incoming and fibril core (odd or even), shift (the alignment of the incoming peptide relative to the template), free chain length (FCL, the number of dissociated residues of the incoming peptide starting from the N- or C-terminus), and the number of hydrogen bonds (Fig. 2 A, B). Simulations were performed to sample the transitions between these registered states as well as transitions between registered and non-registered states (Fig. 2 C and D, SI Table 1). The definition of states and simulation system set-up is described in detail in the Methods section. Figure 3 shows the average transition times of antiparallel registries. The figure suggests that average times of H-bond formation are highly correlated with the local secondary structure of the residue on the incoming peptide. When the residue adopts extended (β-like) conformation, the formation times are relatively short (< 1 ns). In contrast, the H-bond formation times are much longer when the incoming residue is in the coil conformation (up to 8 ns). This secondary structure dependence, however, is distinct from the FCL dependence, which arises from the global movement of the peptide backbone between the bonded residue and peptide terminus. For residues in both extended and coil structures, larger FCL leads to longer H-bond formation times. The overall H-bond formation timescales from explicit solvent simulations are similar to those from implicit solvent simulations(27). The H-bond breakage times, in contrast, are ∼ 5 times slower in explicit solvent. In addition, the breakage rates show weaker dependence on FCL and local secondary structure (Fig. 3C and Fig. S2) and appear to be mainly governed by peptide orientation (antiparallel or parallel) and the residue sidechains. Thus, breakage transitions involving different conformations are combined. Following H-bond breakage, the conformation of the newly free residue (extended/coil) is determined from restrained atomistic simulations, in which the probability of adopting the extended conformation after breakage (*P*_*extend*_) is calculated for each residue. In the MSM simulation, a random number is generated every time a hydrogen bond pair is broken, if the number is less than the *P*_*extend*_ of this residue, the newly free residue will be considered to adopt a β-like extended conformation.

**Figure 2.**
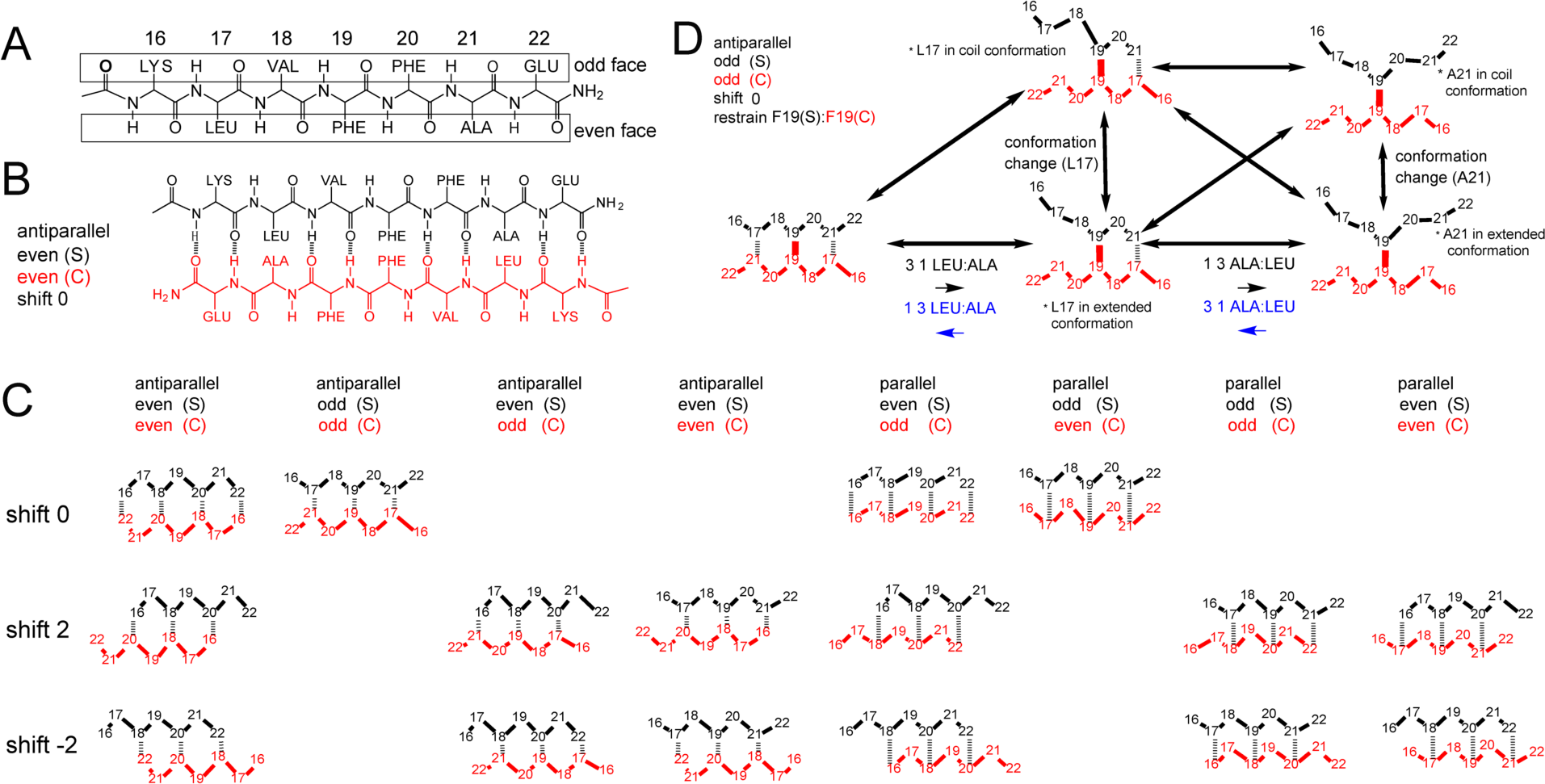
A) Schematic representations of the odd and even faces of a preformed Aβ_16-22_ fibril. B) Schematic representations of H-bonding network of the in-register state “antiparallel e|e|0”. C) all H-bond registries explicitly considered in this work. The odd/even faces and orientation (antiparallel/parallel) of the incoming strand (S) and core peptide (C) are labeled. D) Example transitions of an incoming peptide with the pair 19-19 restrained. Note that the model now includes internal conformational transitions of the incoming peptide (e.g., between “extended” and “coil” states).

**Figure 3.**
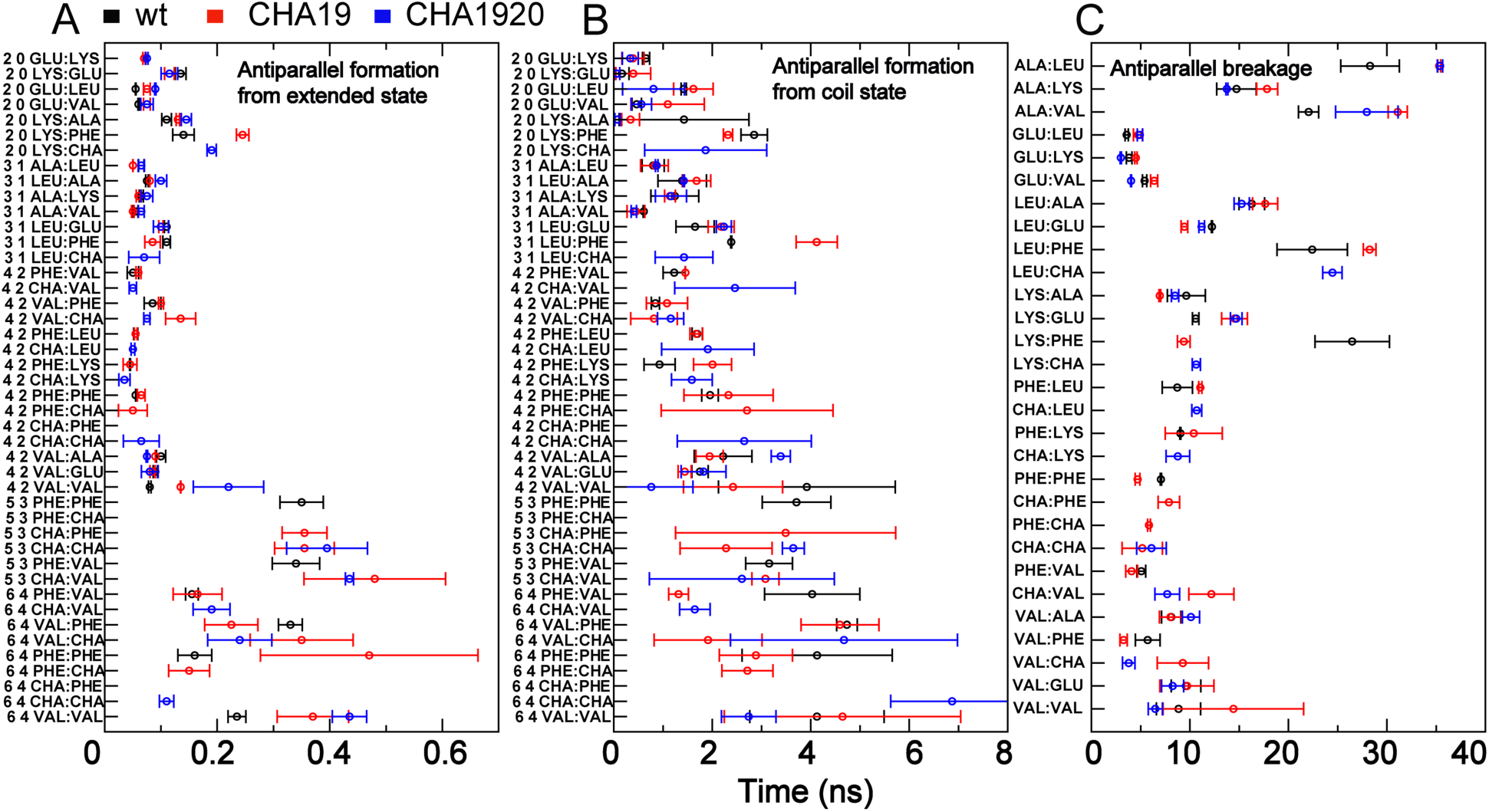
Average backbone H-bond pair transition times associated with anti-parallel registries: A and B) H-bond pair formation from free residue in extended and coil conformation, respectively, C) H-bond pair breakage. All transition times were derived from the restrained simulations where a selected initial H-bond contact pair was harmonically restrained. The pairs are ordered such that those with larger post-transition FCLs are at the bottom.

Similar to our observations in the implicit solvent model, the effects of CHA mutations are not localized to the mutation site; instead, many H-bond transition kinetics are either increased or decreased. Thus, the impact of replacing PHE by CHA on fibril growth could not be directly inferred by changes in H-bond transition rates, but should be attributed to an accumulation of small and mutually compensating effects during the intra- and inter-molecular conformational search during fibril growth.

### Transitions involving non-registered states

Our previous work in implicit solvent highlighted the importance of non-registered states in fibril elongation kinetics. The non-registered states serve as a hub connecting all backbone H-bond registries as well as the disassociated state.(27) In the current study, the non-registered states were further investigated by simulations of the incoming peptide starting from each registry as well as starting from a separation distance *b*. Cluster analysis (see Methods) identified 15 distinct macro-states (Fig. 4). Three of these states are found close to the docking face of the fibril and are connected to registered states. The average transition rates between non-registered states (∼ 1ns) are faster than the average formation rate of the 1^st^ H-bond pair. The transitions between registered states and the three connected non-registered states occur in 2-6 ns. This suggests that the incoming Aβ_16-22_ peptide likely visits many non-registered states before forming the first pair of hydrogen bonds. In addition, for a specific non-registered state, all transitions to registered states can be fitted into a same single-exponential decay curve (Fig S3), which suggest the transition rate between non-specific states and the registered states are independent of orientation (antiparallel or parallel) or residue sidechains, and only depend on the properties of the non-registered state (position, peptide conformation, etc..). Thus, a uniform H-bond formation/breakage transition rate was used for each non-registered state.

**Figure 4.**
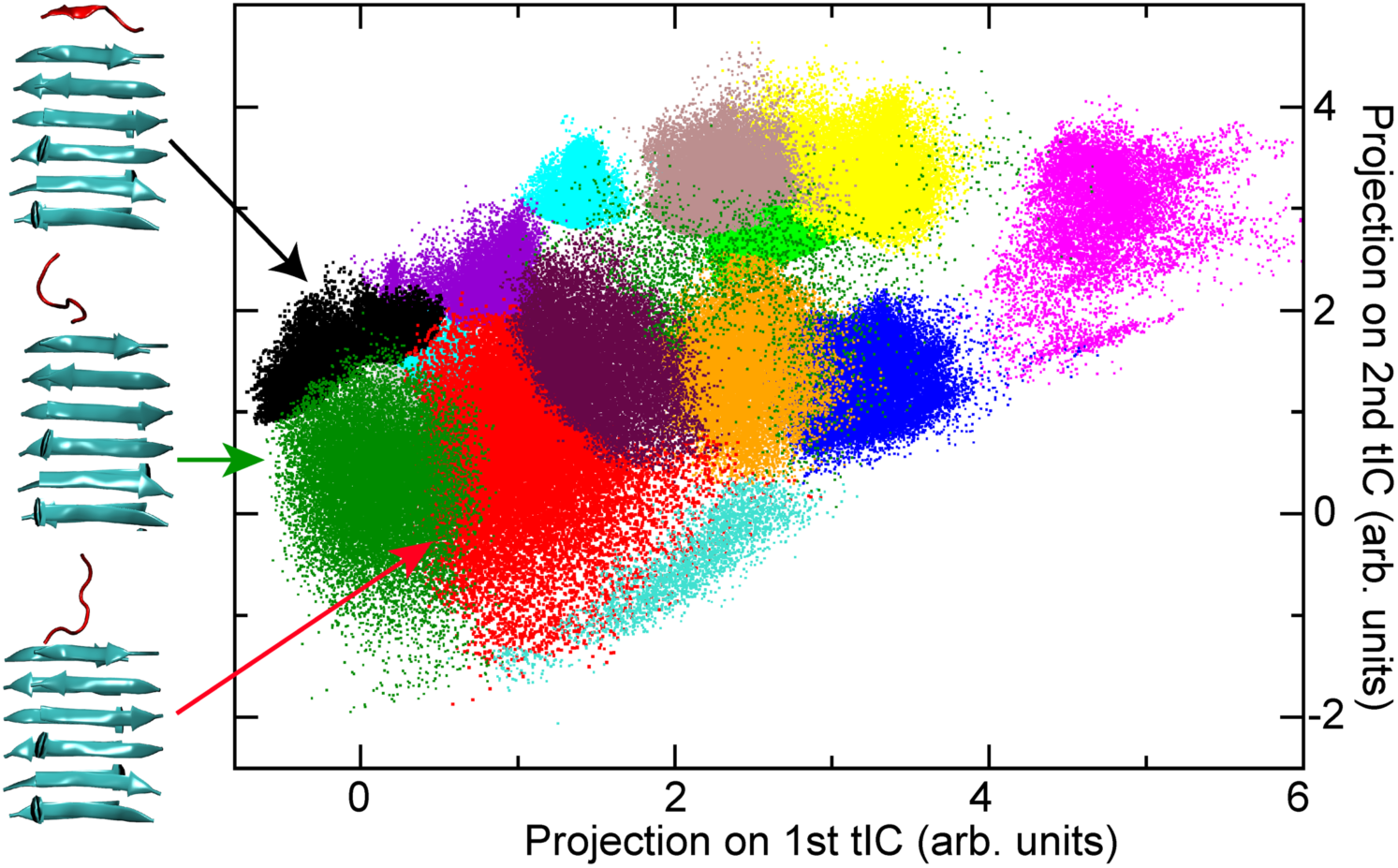
Non-registered state explicitly considered in this work. The simulation trajectories are clustered based on time-lagged independent component analysis (tICA). The states are projected on first two tICAs. The three clusters connected to registered states are colored in black, green and red. Snapshots showing representative conformations the (taken from cluster centers) of the incoming peptide (red) and on the fibril surface (cyan) are also shown. Note, as the cluster is based on 3D tICA, some clusters are covered by other clusters and do not show up in the 2D projection.

### Validation of the MSM model: mutation effects on registry lifetimes

The final MSM model was first examined by comparing the calculated lifetimes of fully H-bonded registries with results derived directly from unrestrained atomistic simulations. Due to the long lifetime in the explicit solvent model, only three short-lifetime antiparallel registries were examined. As shown in Fig. 5, lifetimes predicted by the MSM model (black trace) are essentially identical to results derived from the simulations (solid black dot), suggesting that the MSM model faithfully recapitulates the kinetics (and likely the mechanisms) of the conformational search involved in fibril elongation. This provides a solid basis for applying the final MSM models to further investigate the origin of slow fibril growth kinetics and how it is modulated by mutations.

**Figure 5.**
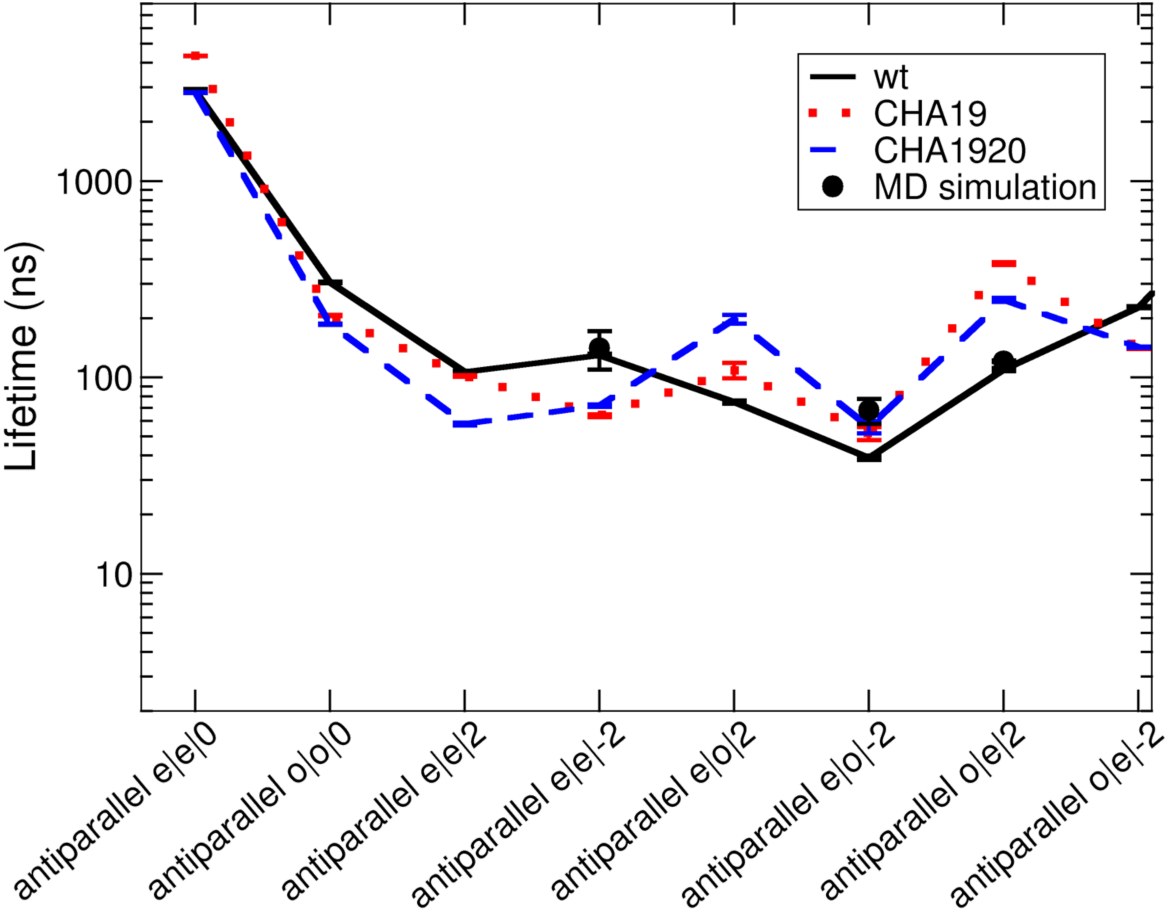
Average lifetimes of fully H-bonded antiparallel registries. Black solid trace and Black dot: MD vs. MSM results for antiparallel registries of the wild-type peptide. Red dot and blue dash traces: MD result of CHA19 and CHA1920 mutated peptides.

We applied the final MSM models to analyze the effects of PHE to CHA mutations on the registry lifetimes. As summarized in Fig. 5, the effects of the single and double CHA mutations are similar on each registry, although the magnitudes of life time changes are different. For both mutations, the time of one in-register state “antiparallel e|e|0” is increased and the life time for the other in-register state “antiparallel o|o|0” is decreased. Also, three antiparallel mis-registered states have longer lifetimes for mutated peptides, and three other mis-registered states have shorter lifetimes. As such, it is not directly obvious what the net effects of mutations have on the overall fibril growth kinetics. Instead, it is necessary to examine the full ensemble of MSM trajectories to account for the ways the peptide may sample various registries before either becoming fully incorporated into the fibril core or returning to solution.

### Molecular origins of slow fibril growth and mutations effects

To derive the overall fibril growth kinetics, 10000 MSM simulations were performed, with 5000 each on the even and odd faces of the fibril core, to calculate *τ*_*residence*_ and *P*_*committor*_ (eq. 1). Each simulation was initiated from the separation distance *b* and terminated when the incoming strand reached the antiparallel fully H-bonded in-register state or the escape distance *q. P*_*committor*_ was calculated as the probability of MSM trajectories reaching the fully H-bonded in-register state, and *τ*_*residence*_ is the average length of all trajectories (Fig, 1B). To calculate *τ*_*off*_, another set of 10000 MSM simulations was initiated from the antiparallel in-register fully bound state and performed until the strand reached the escape distance *q. P*_*committor*_ is nearly doubled compared to our previous study with implicit solvent and without clustering of the non-registered state (Table 1). This suggests the in-register states are more stable for Aβ_16-22_ in the explicit solvent model. In addition, both *τ*_*residence*_ and *τ*_*off*_, are increased by 2∼3 orders of magnitude from the previous study due to the increased residence time of each state (Table 1).

**Table 1.** Key kinetic parameters of fiber growth derived from MSM.

|  | wt | CHA19 | CHA1920 |
| --- | --- | --- | --- |
| $P_{committor}$ | $0.87 \pm 0.08$ | $0.88 \pm 0.09$ | $0.90 \pm 0.02$ |
| $\tau_{residence}$ (ns) | $1359.0 \pm 56.6$ | $546.2 \pm 12.2$ | $588.9 \pm 1.25$ |
| $\tau_{off}$ (ns) | $40012 \pm 515$ | $41733 \pm 836$ | $23086 \pm 522$ |

To compare with experimental results, the elongation rate as a function of monomer concentration is estimated. The diffusion time (*τ*_*diff*_, eq. 1) is estimated as *τ*_*diff*_ = 1/ *k*_*coll*_ *C*_*monomer*_, in which *C*_*monomer*_ is concentration of peptide monomer and *k*_*coll*_ is collision rate constant. The *k*_*coll*_ is defined as *k*_*coll*_*= 4πσD*_*monomer*_, in which the in which *σ* is the reaction cross section (*b* surface in our simulation, 28 Å) and *D*_*monomer*_ is the diffusion coefficient (determined from experiments to be 1.7×10^-6^ cm^2^/s)(14). The results plotted in Fig. 6 suggest the critical concentration of peptide monomer (*C*_*c*_) from the MSM model is ∼8 µM, which is close to the experimental result of ∼33 µM(40). The ability of the current multi-scale modeling approach to recapitulate the critical concentration to a factor of ∼4 without adjustable parameters is noteworthy, particularly considering the comprehensive description of nonspecific and specific reversible binding and the conformational search process. This also likely reflects improvements in explicit solvent force field quality made in recent years.(52, 53)

**Figure 6.**
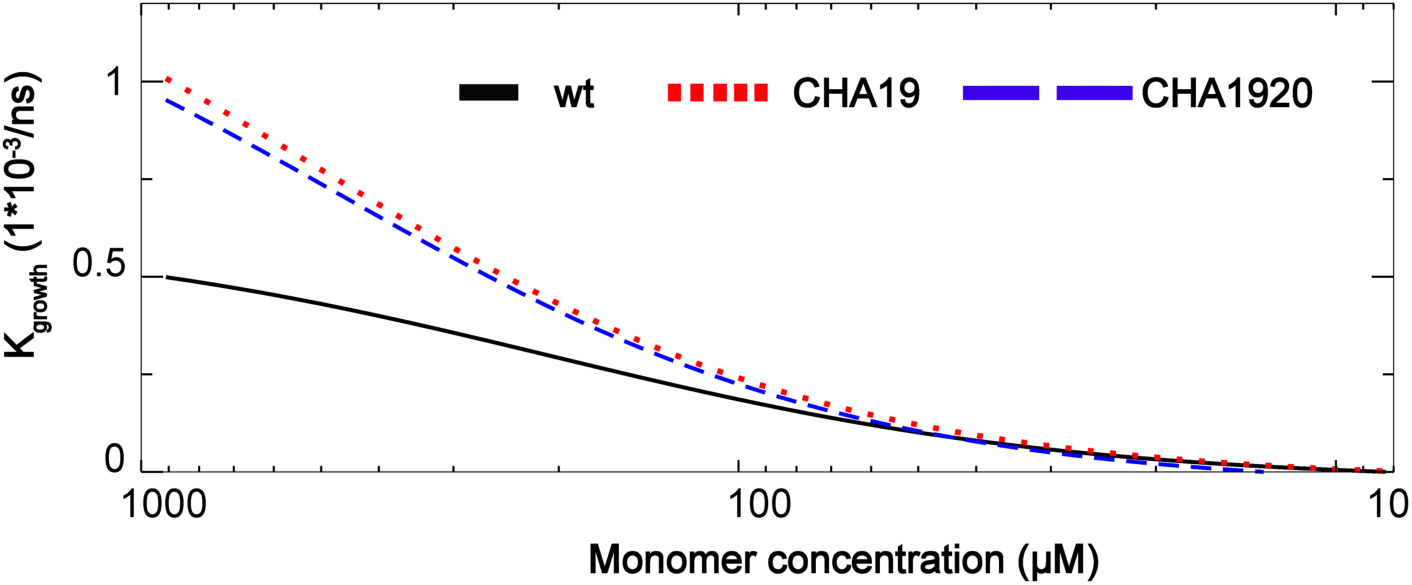
Predicted net growth rate of Aβ_16-22_ fibril as a function of the peptide concentration. The wild type, CHA19, and CHA1920 mutation are presented as black solid trace, red dotted trace and blue dash trace, respectively.

Experimental studies have suggested that CHA mutation on 19 and 20 positions have non-additive effects on overall aggregation kinetics (the growth rate: wt << CHA19 < CHA1920 < CHA20 under high monomer concentration (> 40 µM). However, this result could not be directly explained by thermodynamic effects of the mutations, which appear to have an additive effect (the ΔΔG with respect to wt for CHA19, CHA20, and CHA1920 are 0.2 kcal.mol^-1^, −0.2 kcal.mol^-1^ and 0.0 kcal.mol^-1^, respectively) (40). To investigate the influence of mutations, MSM simulations of CHA mutants were performed. In the initial high monomer concentration condition, both CHA19 and CHA1920 exhibited significant kinetic enhancement in fibril elongation (Fig. 6), in agreement with experiments. (40) This corresponds to the initial stage (0-2h) characterized by a fast decrease in the monomer concentration. This rapid aggregation reduces the monomer concentration leading to an increase of *τ*_*diff*_, which in turn, leads to a decrease of *k*_*growth*_. After the monomer is further depleted the growth rate approaches zero, which suggests that *k*_*on*_ is close to *k*_*off*_. This is the plateau region where the monomer peptide concentration approaches the critical concentration of peptide monomer (*C*_*c*_). (40) Our results also suggest that the wt and CHA mutated peptides have similar *τ*_*diff*_ when *k*_*growth*_ is 0. This is because the time for peptide rearrangement on the fibril surface is much smaller than the diffusion time (*τ*_*residence*_ << *τ*_*diff*_) in this stage. The elongation is then governed by the diffusion process and the effect of mutation can be neglected. This is consistent with the similar measured *C*_*c*_ for wt, CHA19 and CHA1920 (33, 32, 44 µM, respectively).

To investigate the association/dissociation pathways, all MSM trajectories were combined and the relevant free energy of each H-bonding state was plotted (Fig. 7). In marked contrast to the funnel shaped free energy landscape of native proteins(41, 43), the free energy of Aβ_16-22_ is relatively flat. The free energy difference between the in-register sub-states and misregistered sub-states are < 5 kcal/mol. Although the in-register states are more thermodynamically stable (Fig. 5), to reach these states an incoming Aβ_16-22_ peptide needs to visit different mis-registered states an average of ∼22 times before it forms the fully bound in-registered states. The ratio of mis-registered states in amyloid is much higher than the misfolded trapped states observed in natural proteins.(54, 55)

**Figure 7.**
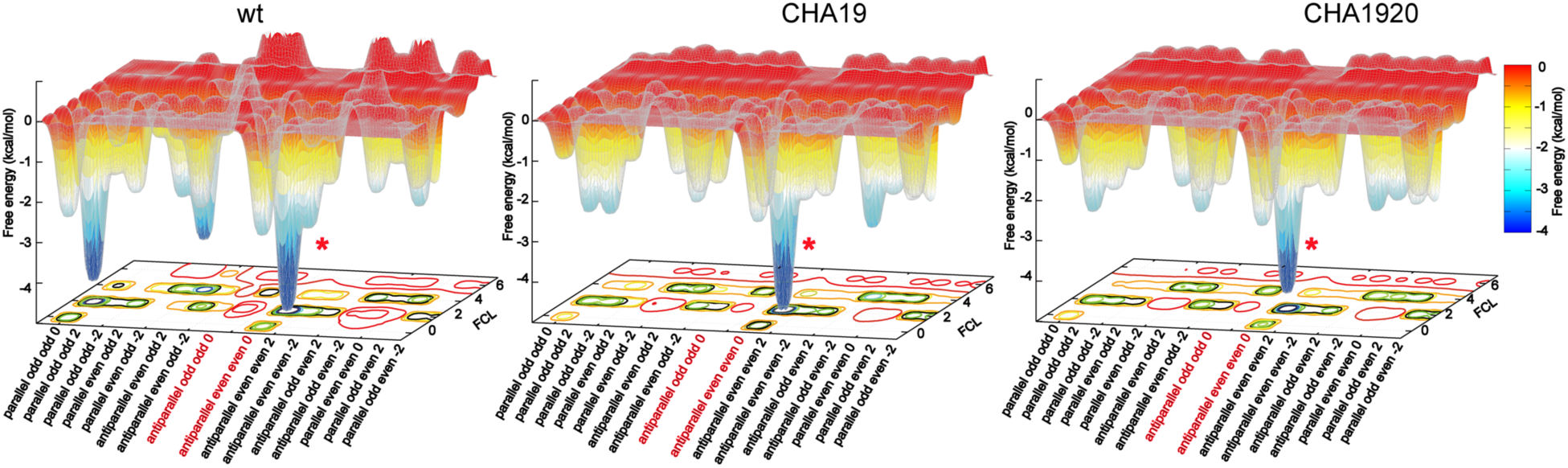
The free energy landscape of Aβ_16-22_ fibril growth. The relative free energy (with respect to “antiparallel e|e|0 FCL 6” sub-state) is derived from all MSM trajectories. The two in-registered states “antiparallel e|e|0” and “antiparallel o|o|0” are highlighted in red. The fully bound “antiparallel e|e|0” state is labelled with a red asterisk.

Our trajectories suggest a molecular interpretation for the “Dock and Lock” model for fibril growth. (12-15). The slow “lock” stage that follows fast nonspecific binding (“dock”) involves a random walk among the non-registered and mis-registered states before the peptide eventually finds the in-register state. The large values of *P*_*committor*_ derived from atomistic simulations (Table 1) show that molecules reaching the docked state are much more likely to proceed to the locked states than to detach. The slow fibril growth rate can be attributed to the time the peptide spends in non-productive trap states. The trapped states, on the other hand, also slow amyloid dissolution by binding the departing peptides in mis-registered states (∼200 times in a single dissociation MSM trajectory). The resulting long *τ*_*off*_ is consistent with the experimental result that the locking stage is nearly irreversible.(12)

To further investigate the influence of mutations, we examine the origin of the change in elongation rate for the mutated peptides. The increased growth rate of the mutants is due to a decrease of *τ*_*residence*_, which in turn, arises from destabilizing two main trap states, “parallel o|o|0” and “parallel e|o|-2” (Table 2). Although the relative free-energy of registered states between the CHA19 and CH1920 double mutation is similar, the double mutation also leads to a decrease in the residence time of in-registered states (Table S2). which leads to a faster dissociation rate (smaller *τ*_*off*_) and a decrease in the growth rate.

**Table 2.** Relative free energy* of registered states in wt or mutated peptides (kcal/mol)

|  | wt | CHA19 | CHA1920 |
| --- | --- | --- | --- |
| antiparallel e\|e\|0 | -11.76 | -8.51 | -7.63 |
| antiparallel o\|o\|0 | -0.72 | -1.14 | -0.32 |
| antiparallel e\|e\|2 | -4.87 | -1.61 | -0.41 |
| antiparallel e\|e\|-2 | -1.86 | -1.04 | -0.63 |
| antiparallel e\|o\|2 | -4.29 | -1.94 | -1.84 |
| antiparallel e\|o\|-2 | 0.00 | -1.02 | -0.48 |
| antiparallel o\|e\|2 | -2.21 | -2.13 | -0.78 |
| antiparallel o\|e\|-2 | -0.17 | -0.52 | -0.21 |
| parallel e\|e\|0 | -3.35 | -0.59 | 0.00 |
| parallel o\|o\|0 | -11.31 | -4.26 | -3.64 |
| parallel o\|o\|2 | -3.21 | -2.65 | -0.88 |
| parallel o\|o\|-2 | -4.32 | -0.33 | -0.22 |
| parallel e\|o\|2 | -3.27 | 0.00 | -0.10 |
| parallel e\|o\|-2 | -6.17 | -0.76 | -0.08 |
| parallel o\|e\|2 | -2.22 | -1.20 | -0.95 |
| parallel o\|e\|-2 | -2.47 | -1.05 | -0.56 |

\* with respect to the state with lowest probability,

### The influence of non-productive pathways

Table 2 shows the free energies for each registry, as estimated from the fraction of time each state was occupied in the MSM. For each molecule the free energies are measured relative to the lowest occupancy registry. In the wt Aβ_16-22_ peptides, two mis-registered states “parallel o|o|0” and “parallel e|o|-2” were found to have similar free energy compared with the in-register states (Table 2). We offer two mechanisms to explain the dominance of the anti-parallel in-register states over these low energy states in the assembled fibril. First, electrostatic interactions will naturally disfavor parallel states due to the close proximity of both terminal and sidechain charges. The repulsion in due to the N and C terminal charges will be reduced somewhat for molecules at the fibril ends because fluctuations in the binding states will allow the charges to separate. However, when the next molecule attaches, those termini will get pinned down, resulting in the full electrostatic repulsion. This effect is amplified with each additional parallel molecule that is added. The electrostatic contribution from the termini becomes less important with increasing protein length, which explains the parallel structures of full-length Aβ, IAPP, and others. In addition, mutations which weaken these repulsions may allow these non-productive pathways to dominate fibril assembly. This assumption is consistent with NMR experiments showing that the E22Q mutation leads to parallel fibrils.(56) Interestingly, this mutation is the same as the Dutch mutant of full-length Aβ peptide which enhances the aggregation propensity.(57)

Secondly, while mis-registered states, like parallel e|o|-2, bind with high affinity, they may offer a poor template for subsequent fibril growth. An out-of-register shift will result in overhanging amino acids that will bind poorly to subsequent molecules due to the conformational entropy penalty of immobilizing the flexible residues.(58) However, under conditions conducive to rapid growth (e.g. when the monomer concentration is high), these registry defects may be trapped within the growing fibril by subsequent monomer addition.(59)

## Conclusion

Functional proteins have evolved to develop efficient folding pathways where a funnel-shaped free energy landscape reduces the time spent exploring non-productive states. This evolutionary pressure does not exist for non-functional states like pathological aggregates. In these cases, the free energy landscape would more closely resemble a golf course (41) or an inverted funnel (59). Without the guidance of a biased landscape, the conformational search will resemble a random search over binding states. This is the situation we find in our simulations. Instead of finding a few dominant pathways or intermediate states, the effects of mutations emerge from the accumulation of small perturbations over the ensemble of partially bound states. This poses new challenges for computational methods, which must sample both productive and non-productive pathways to generate meaningful predictions. We have demonstrated how insights from analytic theory can be utilized to meet this challenge by inspiring sampling techniques that can access new timescales and provide a more complete understanding of fibril growth.

## Methods

A total of 719 µs explicit simulation in the CHARMM36m all-atom force field (60, 61) was performed to derive the microscopic kinetic parameters for transition among all H-bonded and nonspecific bound states. Please see SI Supporting Text for detailed simulation and analysis procedures.

## Acknowledgements

This work was supported by the National Institute of Health (GM107487). Computing for this project was performed on the Beocat Research Cluster at Kansas State University and the Pikes cluster housed in the Massachusetts Green High-Performance Computing Center (MGHPCC).

## References

1. Morriss-Andrews A & Shea J-E (2015) Computational studies of protein aggregation: Methods and applications. Annual Review of Physical Chemistry 66(1):643–666.

2. Fowler DM, Koulov AV, Balch WE, & Kelly JW (2007) Functional amyloid – from bacteria to humans. Trends in Biochemical Sciences 32(5):217–224.

3. Michaels TCT, et al. (2018) Chemical kinetics for bridging molecular mechanisms and macroscopic measurements of amyloid fibril formation. Annual Review of Physical Chemistry 69(1):273–298.

4. Haass C & Selkoe DJ (2007) Soluble protein oligomers in neurodegeneration: lessons from the Alzheimer’s amyloid β-peptide. Nature Reviews Molecular Cell Biology 8(2):101–112.

5. Hardy J & Selkoe DJ (2002) The amyloid hypothesis of Alzheimer’s disease: Progress and problems on the road to therapeutics. Science 297(5580):353–356.

6. Kirkitadze MD, Bitan G, & Teplow DB (2002) Paradigm shifts in Alzheimer’s disease and other neurodegenerative disorders: The emerging role of oligomeric assemblies. Journal of Neuroscience Research 69(5):567–577.

7. Silveira JR, et al. (2005) The most infectious prion protein particles. Nature 437(7056):257–261.

8. Morriss-Andrews A & Shea J-E (2014) Simulations of protein aggregation: Insights from atomistic and Coarse-Grained models. The Journal of Physical Chemistry Letters 5(11):1899–1908.

9. Aisen PS (2005) The development of anti-amyloid therapy for Alzheimer’s disease. CNS Drugs 19(12):989–996.

10. Limbocker R, et al. (2019) Trodusquemine enhances Aβ42 aggregation but suppresses its toxicity by displacing oligomers from cell membranes. Nature Communications 10(1):225.

11. Moreth J, Mavoungou C, & Schindowski K (2013) Passive anti-amyloid immunotherapy in Alzheimer’s disease: What are the most promising targets? Immunity & Ageing 10(1):18.

12. Esler WP, et al. (2000) Alzheimer’s disease amyloid propagation by a template-dependent dock-lock mechanism. Biochemistry 39(21):6288–6295.

13. O’Brien EP, Okamoto Y, Straub JE, Brooks BR, & Thirumalai D (2009) Thermodynamic perspective on the Dock-Lock growth mechanism of amyloid fibrils. The Journal of Physical Chemistry B 113(43):14421–14430.

14. Ban T, Yamaguchi K, & Goto Y (2006) Direct observation of amyloid fibril growth, propagation, and adaptation. Accounts of Chemical Research 39(9):663–670.

15. Collins SR, Douglass A, Vale RD, & Weissman JS (2004) Mechanism of prion propagation: Amyloid growth occurs by monomer addition. PLOS Biology 2(10):e321.

16. Nguyen PH, Li MS, Stock G, Straub JE, & Thirumalai D (2007) Monomer adds to preformed structured oligomers of Aβ-peptides by a two-stage dock–lock mechanism. Proceedings of the National Academy of Sciences 104(1):111–116.

17. Ma B & Nussinov R (2006) Simulations as analytical tools to understand protein aggregation and predict amyloid conformation. Current Opinion in Chemical Biology 10(5):445–452.

18. Straub JE & Thirumalai D (2011) Toward a molecular theory of early and late events in monomer to amyloid fibril formation. Annual Review of Physical Chemistry 62(1):437–463.

19. Wu C & Shea J-E (2011) Coarse-grained models for protein aggregation. Current Opinion in Structural Biology 21(2):209–220.

20. Ban T, et al. (2004) Direct observation of Aβ amyloid fibril growth and inhibition. Journal of Molecular Biology 344(3):757–767.

21. Knowles TPJ, et al. (2007) Kinetics and thermodynamics of amyloid formation from direct measurements of fluctuations in fibril mass. Proceedings of the National Academy of Sciences 104(24):10016–10021.

22. Nguyen HD & Hall CK (2004) Molecular dynamics simulations of spontaneous fibril formation by random-coil peptides. Proceedings of the National Academy of Sciences of the United States of America 101(46):16180–16185.

23. Auer S, Meersman F, Dobson CM, & Vendruscolo M (2008) A generic mechanism of emergence of amyloid protofilaments from disordered oligomeric aggregates. PLOS Computational Biology 4(11):e1000222.

24. Ricchiuto P, Brukhno AV, & Auer S (2012) Protein aggregation: Kinetics versus thermodynamics. The Journal of Physical Chemistry B 116(18):5384–5390.

25. Cheon M, Chang I, & Hall Carol K (2011) Spontaneous formation of twisted Aβ16-22 fibrils in large-scale molecular-dynamics simulations. Biophysical Journal 101(10):2493–2501.

26. Cheon M, Chang I, & Hall CK (2010) Extending the PRIME model for protein aggregation to all 20 amino acids. Proteins: Structure, Function, and Bioinformatics 78(14):2950–2960.

27. Jia Z, Beugelsdijk A, Chen J, & Schmit JD (2017) The Levinthal problem in amyloid aggregation: Sampling of a flat reaction space. The Journal of Physical Chemistry B 121(7):1576–1586.

28. Bacci M, Vymětal J, Mihajlovic M, Caflisch A, & Vitalis A (2017) Amyloid β fibril elongation by monomers involves disorder at the tip. Journal of Chemical Theory and Computation 13(10):5117–5130.

29. Schwierz N, Frost CV, Geissler PL, & Zacharias M (2016) Dynamics of seeded Aβ40-fibril growth from atomistic molecular dynamics simulations: Kinetic trapping and reduced water mobility in the locking step. Journal of the American Chemical Society 138(2):527–539.

30. Han W & Schulten K (2014) Fibril elongation by Aβ17–42: Kinetic network analysis of hybrid–resolution molecular dynamics simulations. Journal of the American Chemical Society 136(35):12450–12460.

31. Schor M, Mey ASJS, Noé F, & MacPhee CE (2015) Shedding light on the Dock–Lock mechanism in amyloid fibril growth using Markov State Models. The Journal of Physical Chemistry Letters 6(6):1076–1081.

32. Schmit JD (2013) Kinetic theory of amyloid fibril templating. The Journal of Chemical Physics 138(18):185102.

33. Barz B, Wales DJ, & Strodel B (2014) A Kinetic approach to the sequence–Aggregation relationship in disease-related protein assembly. The Journal of Physical Chemistry B 118(4):1003–1011.

34. Lane TJ, Shukla D, Beauchamp KA, & Pande VS (2013) To milliseconds and beyond: challenges in the simulation of protein folding. Current Opinion in Structural Biology 23(1):58–65.

35. Senne M, Trendelkamp-Schroer B, Mey ASJS, Schütte C, & Noé F (2012) EMMA: A software package for Markov model building and analysis. Journal of Chemical Theory and Computation 8(7):2223–2238.

36. Pande VS, Beauchamp K, & Bowman GR (2010) Everything you wanted to know about Markov State Models but were afraid to ask. Methods 52(1):99–105.

37. Chodera JD & Noé F (2014) Markov state models of biomolecular conformational dynamics. Current Opinion in Structural Biology 25:135–144.

38. Bowman GR, Huang X, & Pande VS (2009) Using generalized ensemble simulations and Markov state models to identify conformational states. Methods 49(2):197–201.

39. Greenwald J & Riek R (2010) Biology of amyloid: Structure, function, and regulation. Structure 18(10):1244–1260.

40. Senguen FT, et al. (2011) Probing aromatic, hydrophobic, and steric effects on the self-assembly of an amyloid-β fragment peptide. Molecular BioSystems 7(2):486–496.

41. Dill KA & Chan HS (1997) From Levinthal to pathways to funnels. Nature Structural & Molecular Biology 4(1):10–19.

42. Okazaki K-i, Koga N, Takada S, Onuchic JN, & Wolynes PG (2006) Multiple-basin energy landscapes for large-amplitude conformational motions of proteins: Structure-based molecular dynamics simulations. Proceedings of the National Academy of Sciences 103(32):11844–11849.

43. Onuchic JN & Wolynes PG (2004) Theory of protein folding. Current Opinion in Structural Biology 14(1):70–75.

44. Northrup SH, Allison SA, & McCammon JA (1984) Brownian dynamics simulation of diffusion - influenced bimolecular reactions. The Journal of Chemical Physics 80(4):1517–1524.

45. Luty BA, McCammon JA, & Zhou HX (1992) Diffusive reaction rates from Brownian dynamics simulations: Replacing the outer cutoff surface by an analytical treatment. The Journal of Chemical Physics 97(8):5682–5686.

46. Balbach JJ, et al. (2000) Amyloid fibril formation by Aβ16-22, a seven-residue fragment of the Alzheimer’s β-amyloid peptide, and structural characterization by solid state NMR. Biochemistry 39(45):13748–13759.

47. Mehta AK, et al. (2008) Facial symmetry in protein self-assembly. Journal of the American Chemical Society 130(30):9829–9835.

48. Petkova AT, et al. (2004) Solid state NMR reveals a pH-dependent antiparallel β-sheet registry in fibrils formed by a β-amyloid peptide. Journal of Molecular Biology 335(1):247–260.

49. Ferrara P, Apostolakis J, & Caflisch A (2002) Evaluation of a fast implicit solvent model for molecular dynamics simulations. Proteins: Structure, Function, and Bioinformatics 46(1):24–33.

50. Chen J & Brooks CL (2008) Implicit modeling of nonpolar solvation for simulating protein folding and conformational transitions. Phys. Chem. Chem. Phys. 10:471–481.

51. Chen J, Brooks CL, & Khandogin J (2008) Recent advances in implicit solvent based methods for biomolecular simulations. Curr. Opin. Struct. Biol. 18:140–148.

52. Huang J & MacKerell AD (2018) Force field development and simulations of intrinsically disordered proteins. Current Opinion in Structural Biology 48:40–48.

53. Vanommeslaeghe K & MacKerell AD (2015) CHARMM additive and polarizable force fields for biophysics and computer-aided drug design. Biochimica et Biophysica Acta (BBA) - General Subjects 1850(5):861–871.

54. Piana S, Lindorff-Larsen K, & Shaw DE (2013) Atomistic description of the folding of a dimeric protein. The Journal of Physical Chemistry B 117(42):12935–12942.

55. Sborgi L, et al. (2015) Interaction networks in protein folding via atomic-resolution experiments and long-time-scale molecular dynamics simulations. Journal of the American Chemical Society 137(20):6506–6516.

56. Liang C, et al. (2014) Kinetic intermediates in amyloid assembly. Journal of the American Chemical Society 136(43):15146–15149.

57. van Duinen SG, et al. (1987) Hereditary cerebral hemorrhage with amyloidosis in patients of Dutch origin is related to Alzheimer disease. Proceedings of the National Academy of Sciences 84(16):5991–5994.

58. Zhang L & Schmit JD (2016) Pseudo-one-dimensional nucleation in dilute polymer solutions. Physical Review E 93(6):060401.

59. Huang C, Ghanati E, & Schmit JD (2018) Theory of Sequence Effects in Amyloid Aggregation. The Journal of Physical Chemistry B 122(21):5567–5578.

60. Huang J & MacKerell AD (2013) CHARMM36 all-atom additive protein force field: Validation based on comparison to NMR data. Journal of Computational Chemistry 34(25):2135–2145.

61. Huang J, et al. (2016) CHARMM36m: an improved force field for folded and intrinsically disordered proteins. Nature Methods 14:71.

62. Jorgensen WL, Chandrasekhar J, Madura JD, Impey RW, & Klein ML (1983) Comparison of simple potential functions for simulating liquid water. The Journal of Chemical Physics 79(2):926–935.

