## Supplementary material for "The dark side of amyloid aggregation: Exploring the productive and non-productive pathways with multi-scale modeling": Methods and supplemental data

### MSM model for prediction the amyloid aggregation

### Methods

#### Enumeration of peptide H-bond registries

To enumerate all possible register states, we first define the registries by the peptide orientation (antiparallel or parallel), contact surface ("even" or "odd") and alignment (shift -2, 0 or 2) using a notation similar to our previous work(1) (Fig. 2 A and B). As illustrated in Fig. 2 B, an in-register state "antiparallel e|e|0" denotes the state where the even face of incoming peptide docks on the even face of the fibril core in antiparallel orientation with 0 shift (in-register). As the two H-bonds involving the same residue are close in space and are highly correlated in both formation and breakage transitions(1) (Fig. 2A), the transitions in the MSM model present the formation and breakage of H-bond pairs rather than individual bonds. The shift value increases or decreases by 2 between registries docked with same surfaces. Note that, although registries with up to 6/-6 are possible, the binding lifetimes scale exponentially with the number of possible H-bonds(2, 3). As such, states with two or fewer pairs of H-bonds contribute minimally to the sampling of registries. These states are thus grouped into the non-registered states and not described explicitly in the MSM model. In the current model, the hydrogen bond pair is identified by the peptide orientation and the pair of incoming/core peptide residues involved. For example, there are four possible H-bond pairs in the "antiparallel e|e|0" state (antiparallel 16-22, antiparallel 18-20, antiparallel 20-18 and antiparallel 22-16); the "antiparallel e|e|0" registry contains a series of sub-states defined by the formed hydrogen bond pair (e.g. 16-22|18-20 represent the states which the two hydrogen bond pairs on the N-terminus of incoming peptide is formed, whereas the other residues in the C-terminus are still unbound and free).

#### Explicit solvent sampling of transitions between H-bond states

All simulations were performed using Gromacs2016(4, 5) with the CHARMM36m all-atom force field(6, 7) and TIP3P water model(8). The MD time step was set at 2 fs. Electrostatic interactions were described by using the Particle Mesh Ewald (PME) algorithm(9) with a cutoff of 12 Å. Van der Waals interactions were cutoff at 12 Å with a smooth switching function starting at 10 Å. Covalent bonds to hydrogen atoms were constrained by the SHAKE algorithm.(10) The temperature was maintained at 298 K using the Nose-Hoover thermostat(11, 12). The pressure was maintained semi-isotopically at 1 bar using the Parrinello–Rahman barostat algorithm.(13)

In all simulations, the positions of all backbone heavy atoms in the core were harmonically restrained using a force constant of 10 kcal/mol/Å$^2$. The solvated systems were then neutralized by adding 150 mM NaCl.

It has been shown that A$\beta_{16\text{-}22}$ forms bilayer or multilayer β-sheets in solution.(14, 15) As such, we included two layers of anti-parallel β-sheets to represent the preformed fibril. Specifically, as illustrated in Fig. S1, each layer contains a pentamer, which corresponds to the approximate size of the critical nucleus of fibril formation.(16, 17) Each fibril has two docking sites available for incoming peptides (Fig. S1, yellow strand on each end of the fibril). Therefore, for each registry two incoming peptides were first docked on the core in fully H-bonded conformations (one on each end). Then for each hydrogen bond pair within the registry a set of 50 production simulations of 50 ns each in length were performed with a single hydrogen bond pair restrained (e,g, Fig 2C shows the restraint of bond 19-19). The aim of these simulation is sampling of H-bond transitions around the anchoring pair (e.g. breakage and formation of 17-21 and 21-17 when 19-19 is restrained). Before each simulation starts, the heavy atoms of the incoming residue involved in the H-bond pair are restrained and the system is heated at 800 K for 100 ps. During heating, the unrestrained part of incoming peptide is partially disordered (the core peptides will not be affected as they are always restrained). Thus, each simulation starts from a random state in which 1-4 H-bond pairs are formed

In the previous implicit solvent study(1), it was found that the kinetics of H-bond transitions mainly depend on the orientation of the peptide (parallel vs. antiparallel), the contacting residues, and the length of “free” peptide chains adjacent to the given H-bond pair before/after transition (free chain length, or FCL). For example, the transition “antiparallel 2 0 LYS:GLU” describes the formation of an antiparallel hydrogen bond pair of the terminal residues. The formation of H-bonds between LYS and GLU on the incoming and core peptides, respectively, results in a reduction in the FCL from two to zero. The implication is that the peptide reconfiguration time is fast in implicit solvent and H-bond transition kinetics are independent of the dissociated peptide conformational state. However, this is no longer true when explicit solvent is included in simulation. A$\beta_{16\text{-}22}$ is short and intrinsically disordered; we observed that the formation kinetics of a given H-bond pair, as noted above, also depends on the secondary structure state of the incoming peptide. Furthermore, analysis of the transition times shows that only two states, "extended" and "coil", need to be considered. For a particular residue on the incoming peptide, the state is defined by the secondary structure of the preceding and/or subsequent residues (if available). For example, for residue 17, the secondary structure of 16-18 is first determined by their phi/psi angle (Fig S4). If residues 16-18 all adopt β structure, the residue 17 is considered to be in the "extended" states. If any of these residues does not adopt β structure, then residue 17 is assigned to the "coiled" state. As such, two sets of formation kinetic parameters will be collected for each H-bond pair, depending on whether the residue of the incoming peptide is in "extended" or "coil" state. Accordingly, the corresponding H-bond sub-states from our previous MSM model are split into two states, depending on the secondary structure state of the incoming peptide (see Fig. 2D).

**Simulations and cluster analysis of non-registered states**

To simulate the transitions involving non-registered states, two sets of unrestrained simulations were performed. The first set of simulations focused on the transition between non-registered states and registered states. These simulations were started using the same structure as the restrained simulations described above, but with only the backbone heavy atoms of the fibril core harmonically restrained. The incoming peptides were unrestrained and could freely sample various bound and unbound states (Fig. 2D, Table S1). The second set of simulations are performed to examine the diffusion-collision rates of the incoming peptides with the fibril core. These simulations focus on sampling the transitions between non-registered states, the separation distance *b*, and the escape distance *q*. Peptides were initially placed in a random position 28 Å from the core layer (centers of mass separation). This distance is chosen as it is close to half of the end-to-end distance of amyloid 16-22 peptide in a fully extended state, plus the nonbond cutoff distance of the simulation (12 Å in this work). This distance is sufficiently large such that the inter-molecular potential of mean force is only a function of the separation between the incoming peptide and core peptides and not their relative orientation.(18, 19) The *q* distance is set to 2*b* (56 Å). A relatively large simulation box (120 * 120 * 120 Å) is used in these simulations, A total of 20 trials of 200 ns each in length were performed (see Fig 2D).

Clustering analysis was applied to identify conformational sub-states involved in the nonspecific bound states. Three classes of features are chosen for clustering analysis. The first set of features, hydrogen bond states and SASA contact areas, describe the nature of contacts between incoming peptides and the fibril core. The second set of features, incoming peptide end-to-end distance and the number of residues in the extended (beta) conformation, describe the peptide internal conformation. The last set of features represent the relative positioning between incoming peptides and fibril core, including center of mass (COM) distance between the incoming peptide and the docking layer of the core, minimum distance between the incoming peptide and top layer of the core, and minimum distance between the incoming peptide and the rest of the core.

These features are normalized to have close to zero mean and unit variance using a Standard Scaler method(20, 21). Then the time-lagged independent component analysis (tICA) method is used to reduce the dimensionality.(22-24) The simulation trajectories are clustered into micro-states using a hybrid k-centers k-medoids algorithm(21). The kinetically related micro-states are lumped into macro-states using Perron Cluster Cluster Analysis (PCCAplus) algorithm(25, 26). The generalized matrix Rayleigh quotient (GMRQ) method(24, 27) is applied to optimize the hyperparameters as described in previous studies(28, 29).

**Simulations of mutated peptides and model validation**

The first set of simulations (the incoming peptide is restrained in all sub-states for each registry) (see Fig 2D) were repeated for two mutant sequences in which phenylalanine residues at positions 19 or 19/20 are replaced with the non-natural amino acid cyclohexylalanine (CHA19 and CHA1920). For the second set of simulations (the incoming peptide is unrestrained in all sub-states of the registry), only the registries directly related to the mutation (e.g. antiparallel o|o|0 which contains three pair 17-21, 19-19 and 21-17) were repeated. To validate the MSM model, an additional set of simulations was performed for two short lifetime antiparallel registries of the wild type peptide, where 100 simulations were initiated from fully H-bonded conformations lasting 200 ns each (Fig 2D). These simulations yield the lifetimes of each registry in the fully bound state and provide a direct validation of the lifetimes predicted by the MSM model.

**Markov state modeling of fibril growth**

Two sets of MSM simulations were performed. In the first set of MSM simulations, the peptide was started from the *b* surface and the simulations continued until the peptide evolved to either the in-registered fully bound (antiparallel o|o|0 or antiparallel e|e|0) or dissociated states (*q* surface), From these simulations, $\tau_{residence}$ and $P_{committor}$ were derived. The second set of simulations started from fully bound in-registered states, and continued until the dissociated state was reached. From these simulations $\tau_{off}$ is derived. Each set of simulations was performed 10000 times

(5000 for fibril cores with as even surface docking layer and 5000 for cores with an odd surface docking layer). The Gillespie algorithm(30) was employed to generate stochastic trajectories of fibril growth. For each step, two random numbers ($R_1$ and $R_2$) in the interval [0, 1] are generated. Given the rates $k_1, k_2, \ldots, k_n$ for all possible transitions from the current state and the sum of these rates, $k_{tot}$, transition $i+1$ is selected when

$$\frac{k_i}{k_{tot}} < R_1 < \frac{k_{i+1}}{k_{tot}} \tag{2}$$

The elapsed time required before transition $i+1$ occurs is set equal to

$$t = -\frac{1}{k_{tot}} ln(1 - R_2) \tag{3}$$

The chosen state and elapsed time are appended to the trajectory, at which point the new set of accessible states is determined and the algorithm repeats.

**Table S1 Summary of atomistic production simulations**

| Purpose | H-bond transitions around the anchoring pair | Transition between H-bonded and non-registered states | Transition between the *b*, q surfaces and non-registered states | Lifetimes of fully bound registries states |
|---|---|---|---|---|
| Peptides | WT, CHA19, and CHA1920 | WT, CHA19, and CHA1920 | WT | WT |
| Initial structures | Singly H-bonded register states | Singly H-bonded register states | random position on b surface | Fully H-bonded register state |
| Restraints | Fibril core and the initial H-bond contact pair | Fibril core | Fibril core | Fibril core |
| Simulations | 50 ns × 50 (runs) × 50 (register states) (for all three peptides) | WT: 50 ns × 50 (runs) × 50 (register states)<br>CHA19: 50 ns × 50 (runs) × 30 (CHA19 perturbed register states)<br>CHA20: 50 ns × 50 (runs) × 44 (CHA1920 perturbed register states) | 200 ns × 20 (runs) | 200 ns × 50 (runs) × 3 (antiparallel registries) |
| Box size (Å) | 80 x 60 x 60 | 80 x 60 x 60 | 120 x 120 x 120 | 80 x 60 x 60 |

**Table S2 Averaged residence time* of registered states in wt or mutated peptides (ns)**

| | wt | CHA19 | CHA1920 |
|---|---|---|---|
| antiparallel e\|e\|0 | 83.8 | 66.59 | 36.77 |
| antiparallel o\|o\|0 | 0.45 | 1.86 | 1.6 |
| antiparallel e\|e\|2 | 2.65 | 0.72 | 0.45 |
| antiparallel e\|e\|-2 | 0.41 | 0.56 | 0.62 |
| antiparallel e\|o\|2 | 1.44 | 0.94 | 1.51 |
| antiparallel e\|o\|-2 | 0.07 | 0.42 | 0.49 |
| antiparallel o\|e\|2 | 1.03 | 3.61 | 1.99 |
| antiparallel o\|e\|-2 | 0.29 | 0.88 | 1.23 |
| parallel e\|e\|0 | 1.68 | 0.96 | 0.98 |
| parallel o\|o\|0 | 35.72 | 1.73 | 1.89 |
| parallel o\|o\|2 | 0.44 | 1.77 | 0.73 |
| parallel o\|o\|-2 | 0.96 | 0.21 | 0.33 |
| parallel e\|o\|2 | 1.79 | 0.56 | 1.05 |
| parallel e\|o\|-2 | 10.22 | 1.15 | 1.2 |
| parallel o\|e\|2 | 0.42 | 0.57 | 0.75 |
| parallel o\|e\|-2 | 0.35 | 0.57 | 0.64 |

* Note, the average residence time include sub-registered states with different FCL length, which is different from the lifetime of the states start from H-bonded antiparallel registries (Fig. 6)

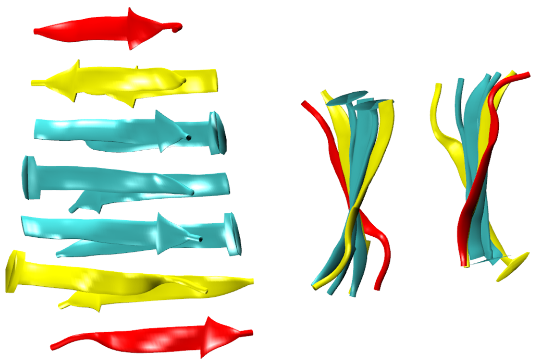

**Figure S1**. Side (left) and top views (right) of a representative initial structure of explicit solvent simulations (antiparallel e|e|0 registry). The peptides are shown in cartoon representations. The docking faces of the fibril core are colored in yellow and incoming peptides in purple.

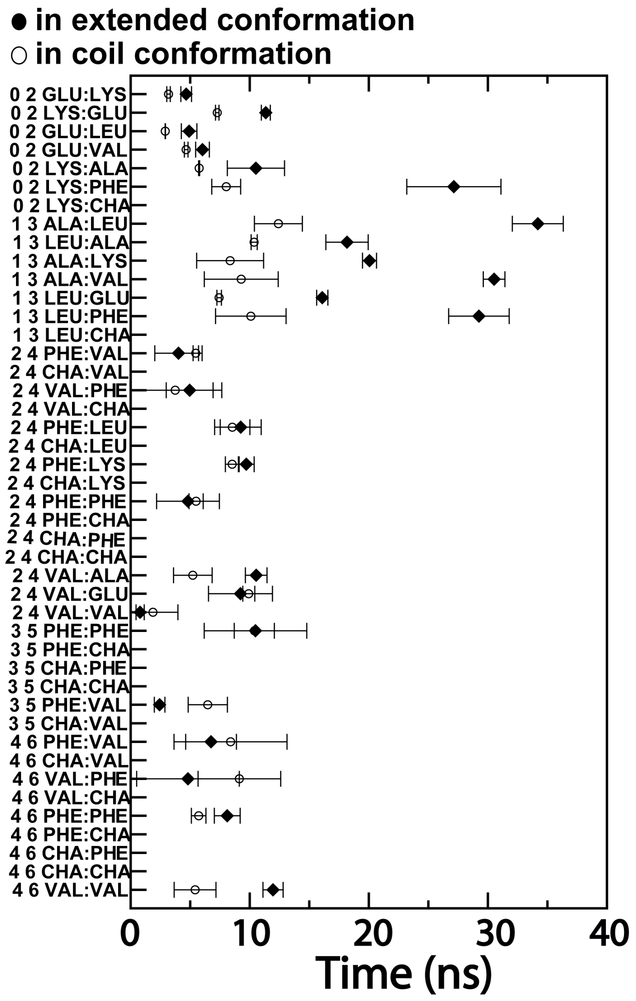


**Figure S2.** Average backbone H-bond pair breakage times involved different local secondary states. The solid dot represent transitions when new free residue adopt extended conformation, the open dot represent transitions when new free residue adopt coil conformation.

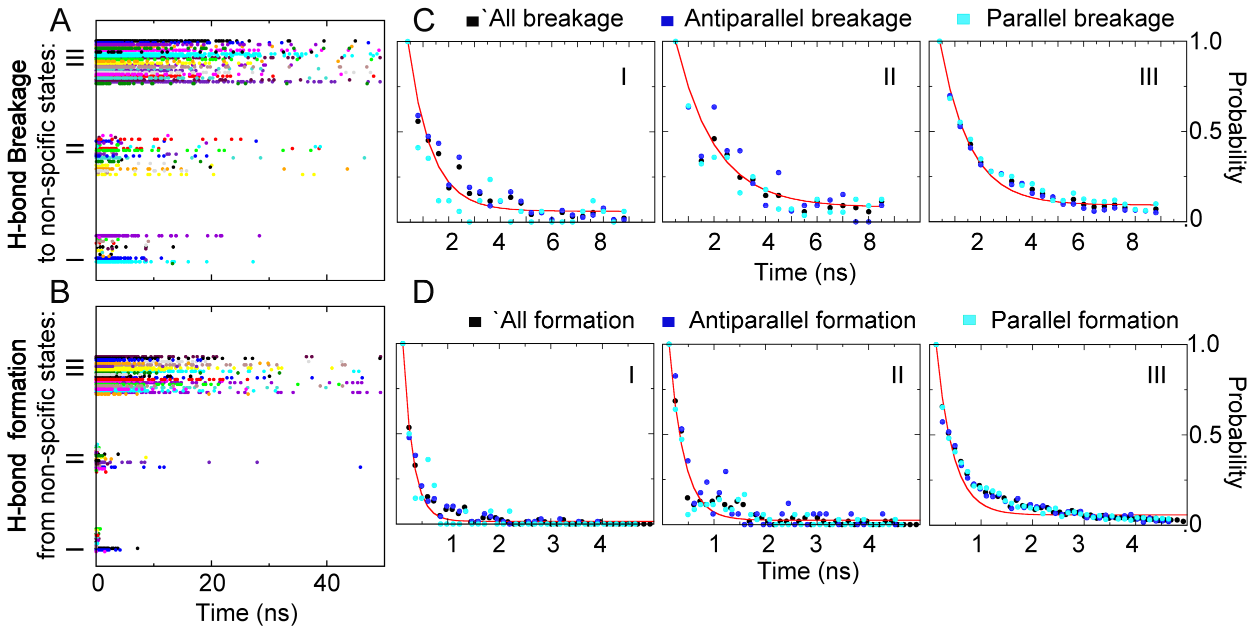


**Figure S3.** Transitions between registered states and non-registered states. There are three non-registered states connect to registered states, they are labelled as state I, II and III. Left panel: Transition event observed in simulations. Each individual transition event are represented as a dot in the figure. The residue pair/sidechain type are indicated by colors. Right panel: The fitted curve (red trace) use the histogram from all events (black dot). The histogram of transitions involved antiparallel orientation (blue dot) or parallel orientation (cyan dot) adopt a similar distribution around the curve.

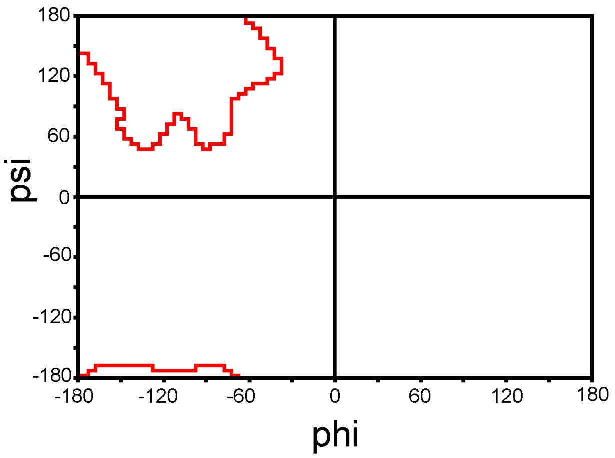


**Figure S4.** The Ramachandran plot of the beta region (inside red traces) in peptide secondary structure analysis.

#### References


1. Jia Z, Beugelsdijk A, Chen J, & Schmit JD (2017) The Levinthal problem in amyloid aggregation: Sampling of a flat reaction space. *The Journal of Physical Chemistry B* 121(7):1576-1586.
2. Schmit JD (2013) Kinetic theory of amyloid fibril templating. *The Journal of Chemical Physics* 138(18):185102.
3. Huang C, Ghanati E, & Schmit JD (2018) Theory of Sequence Effects in Amyloid Aggregation. *The Journal of Physical Chemistry B* 122(21):5567-5578.
4. Hess B, Kutzner C, van der Spoel D, & Lindahl E (2008) GROMACS 4: algorithms for highly efficient, load-balanced, and scalable molecular simulation. *Journal of Chemical Theory and Computation* 4(3):435-447.
5. Abraham MJ*, et al.* (2015) GROMACS: High performance molecular simulations through multi-level parallelism from laptops to supercomputers. *SoftwareX* 1–2:19-25.
6. Huang J & MacKerell AD (2013) CHARMM36 all-atom additive protein force field: Validation based on comparison to NMR data. *Journal of Computational Chemistry* 34(25):2135-2145.
7. Huang J*, et al.* (2016) CHARMM36m: an improved force field for folded and intrinsically disordered proteins. *Nature Methods* 14:71.
8. Jorgensen WL, Chandrasekhar J, Madura JD, Impey RW, & Klein ML (1983) Comparison of simple potential functions for simulating liquid water. *The Journal of Chemical Physics* 79(2):926-935.
9. Darden T, York D, & Pedersen L (1993) Particle mesh Ewald: An *N*-log (*N*) method for Ewald sums in large systems. *The Journal of Chemical Physics* 98:10089.
10. Ryckaert J-P, Ciccotti G, & Berendsen HJC (1977) Numerical integration of the cartesian equations of motion of a system with constraints: molecular dynamics of n-alkanes. *Journal of Computational Physics* 23(3):327-341.
11. Hoover WG (1985) Canonical dynamics: Equilibrium phase-space distributions. *Physical Review A* 31(3):1695-1697.
12. Nosé S (1984) A unified formulation of the constant temperature molecular dynamics methods. *The Journal of Chemical Physics* 81(1):511-519.
13. Parrinello M & Rahman A (1981) Polymorphic transitions in single crystals: A new molecular dynamics method. *Journal of Applied Physics* 52(12):7182-7190.
14. Cheon M, Chang I, & Hall Carol K (2011) Spontaneous formation of twisted Aβ16-22 fibrils in large-scale molecular-dynamics simulations. *Biophysical Journal* 101(10):2493-2501.
15. Senguen FT*, et al.* (2011) Probing aromatic, hydrophobic, and steric effects on the self-assembly of an amyloid-β fragment peptide. *Molecular BioSystems* 7(2):486-496.
16. Goldsbury CS*, et al.* (2000) Studies on the in vitro assembly of Aβ 1–40: Implications for the search for Aβ fibril formation inhibitors. *Journal of Structural Biology* 130(2):217-231.
17. Hills RD & Brooks CL (2007) Hydrophobic cooperativity as a mechanism for amyloid nucleation. *J. Mol. Biol.* 368(3):894-901.
18. Northrup SH, Allison SA, & McCammon JA (1984) Brownian dynamics simulation of diffusion ‐ influenced bimolecular reactions. *The Journal of Chemical Physics* 80(4):1517-1524.

19. Luty BA, McCammon JA, & Zhou HX (1992) Diffusive reaction rates from Brownian dynamics simulations: Replacing the outer cutoff surface by an analytical treatment. *The Journal of Chemical Physics* 97(8):5682-5686.
20. Pedregosa F, *et al.* (2011) Scikit-learn: Machine learning in Python. *Journal of machine learning research* 12(Oct):2825-2830.
21. Beauchamp KA, *et al.* (2011) MSMBuilder2: Modeling conformational dynamics on the picosecond to millisecond scale. *Journal of Chemical Theory and Computation* 7(10):3412-3419.
22. Deuflhard P, Huisinga W, Fischer A, & Schütte C (2000) Identification of almost invariant aggregates in reversible nearly uncoupled Markov chains. *Linear Algebra and its Applications* 315(1):39-59.
23. Schwantes CR & Pande VS (2013) Improvements in Markov state model construction reveal many non-native interactions in the folding of NTL9. *Journal of Chemical Theory and Computation* 9(4):2000-2009.
24. McGibbon RT & Pande VS (2015) Variational cross-validation of slow dynamical modes in molecular kinetics. *The Journal of Chemical Physics* 142(12):124105.
25. McGibbon RT, Husic BE, & Pande VS (2017) Identification of simple reaction coordinates from complex dynamics. *The Journal of Chemical Physics* 146(4):044109.
26. Deuflhard P & Weber M (2005) Robust Perron cluster analysis in conformation dynamics. *Linear Algebra and its Applications* 398:161-184.
27. Husic BE, McGibbon RT, Sultan MM, & Pande VS (2016) Optimized parameter selection reveals trends in Markov state models for protein folding. *The Journal of Chemical Physics* 145(19):194103.
28. Sultan MM, Denny RA, Unwalla R, Lovering F, & Pande VS (2017) Millisecond dynamics of BTK reveal kinome-wide conformational plasticity within the apo kinase domain. *Scientific Reports* 7(1):15604.
29. Jayachandran G, Vishal V, & Pande VS (2006) Using massively parallel simulation and Markovian models to study protein folding: Examining the dynamics of the villin headpiece. *The Journal of Chemical Physics* 124(16):164902.
30. Gillespie DT (1977) Exact stochastic simulation of coupled chemical reactions. *The Journal of Physical Chemistry* 81(25):2340-2361.
